# A new player in the biochemistry of Anammox bacteria: a multidomain HAO-like protein

**DOI:** 10.64898/2026.07.28.741245

**Authors:** Susana F. Fernandes, Catarina Martins Alves, Catarina M. Paquete, Ricardo O. Louro, Filipe Folgosa

**Affiliations:** Instituto de Tecnologia Química e Biológica António Xavier, Universidade Nova de Lisboa, Av. da República, 2780-157 Oeiras, Portugal

**Keywords:** Multicopper Oxidase, Hydroxylamine Oxidoreductases, Anammox cycle, Hydroxylamine reduction, Multi-domain enzymes, Hydrazine production

## Abstract

Anaerobic ammonium-oxidizing (anammox) bacteria are essential players in the global nitrogen cycle, responsible for converting ammonium and nitrite directly to nitrogen gas. Anammox bacteria have unique features such as a specialized cellular compartment - the anammoxosome. *Candidatus (Ca.) Brocadia pituitae* genome, as other anammox bacteria, encodes for a diversity of hydroxylamine oxidoreductase (HAO) paralogs, often pointed out as the enzymes responsible for most of the reactions of the anammox cycle. One of this *Ca. B. pituitae* HAO paralogs is an 840-amino acids protein, named here as BpMHAO, that stands out for its unprecedented domain organization, which includes a multicopper oxidase-like (MCo-like) domain followed by the HAO-like one. Sequence and structural analyses classified this MCo-like domain as homologous to the small laccase family. Spectroscopic characterization revealed a distinct UV-visible spectrum, tentatively assigned to the T3 center, whereas the EPR spectra confirmed the presence of T1, T2 and T3 copper centers. Enzymatic studies demonstrated limited laccase and oxygen-dependent ferroxidase activities. On the other hand, enzymatic assays performed in cell extracts from *Escherichia coli* and *Shewanella oneidensis*, harbouring the recombinant HAO-like domain, exhibited a robust hydroxylamine reductase activity using methyl viologen as the electron donor. Our results showed that the BpMHAO potentially plays a role in the anammox process/reactions by converting hydroxylamine into hydrazine. This feature can be relevant to anammox bacteria either by i) mitigating unwanted hydroxylamine, obtained by incorrect formation of this compound, by converting it into hydrazine and enabling its use in the anammox reaction or ii) using hydroxylamine from the outside medium as a substitute for ammonium, delivering hydrazine directly to the last step of the cycle, short-circuiting its first steps.

## Introduction

Ammonium (NH_4_^+^) accumulation in aquatic environments poses serious ecological and public health risks and remains a central challenge for wastewater treatment. Conventional biological nitrogen removal relies on sequential nitrification and denitrification, typically requiring high aeration inputs and sufficient organic carbon, while producing substantial sludge and greenhouse gas emissions [1]. These constraints call for more energy- and resource-efficient nitrogen removal solutions.

Anaerobic ammonium oxidation (anammox) is a microbially mediated process where ammonium is oxidized with nitrite (NO₂⁻) as electron acceptor under anoxic conditions, producing dinitrogen gas (N_2_), with the formation of nitric oxide (NO) and hydrazine (N_2_H_4_) as intermediates, in an unique cell compartment, the anammoxosome [2,3]. Since its discovery, anammox bacteria became a cornerstone technology for sustainable nitrogen removal due to reduced oxygen demand, lower sludge production and potentially lower carbon emissions compared with conventional approaches [4–6]. The chemolithoautotrophic anammox bacteria belong to the phylum Planctomycetota, with key genera including *Candidatus Brocadia*, *Candidatus Kuenenia*, *Candidatus Scalindua* and *Candidatus Jettenia* [7,8]. The majority of the anammox reactions were proposed to be performed by hydroxylamine oxidoreductases (HAO)-like proteins, implicating their ability to perform a wide range of activities, such as nitrite reduction, hydroxylamine oxidation and hydrazine dehydrogenase [9–12]. It was hypothesized that the anammoxosome can harbour up to 12 homologous of HAO-like proteins [13].

Hydroxylamine oxidoreductases are characterized by a complex, multi-heme architecture, generally organized as homotrimers, where each monomer contains eight covalently bound *c*-type hemes, resulting in a total of 24 hemes per trimeric enzyme [14–16]. Seven of these hemes are standard low-spin *c*-type hemes, with a bis-histidine axial coordination and covalently linked to the protein backbone via two thioether bonds to cysteine residues, whereas the eighth is the unique, catalytic “P_460_ cofactor”, with a third, unique covalent bond between a meso-heme carbon and an aromatic ring carbon on a conserved tyrosine residue [17]. This tyrosine-porphyrin cross-link is thought to modulate the active site by distorting the heme plane, thereby facilitating the abstraction of electrons from the substrate, therefore associated to oxidation reactions [10,18].This heme, usually assigned as the heme-4, the catalytic one, has an hig-spin configuration with a single axial histidine coordination. Research on *K. stuttgartiensis* indicates that the absence of the conserved tyrosine cross-link in its HAO homolog tunes the enzyme toward reductive catalysis, contrasting with the oxidative nature of the nitrifier enzyme [10].

In anammox bacteria, other structural organizations have been observed, such as the supercomplex of hydrazine dehydrogenases from *Kuenenia stuttgartiensis* and *Brocadia fulgida*, containing 192-hemes packed sufficiently close for fast intramolecular electron transfer [19].

The majority of anammox bacteria with known genomes encode for an uncommon gene, *kuste2479* in *Ca. K. stuttgartiensis* [20,21]. The protein codified by this gene is formed by an unique combination of two domains: a multicopper oxidase (MCo)-like domain, homologous to the family of small laccases, and a HAO-like domain [22,23].

Multicopper oxidases (MCo) are a family of copper-containing proteins, widely distributed in nature, and include examples such as ascorbate oxidases, laccases, small laccases, and laccase-ferroxidases, that play a role in the oxidation of various substrates, such as phenolic compounds, aromatic amines, and several inorganic compounds, using oxygen as an electron acceptor [24,25]. Essential to their activity are the four copper ions, organized into three types of centers: T1 (mononuclear “blue type center”), where the oxidation of the substrate occurs; T2 and T3 (binuclear copper center), which form a trinuclear center (TNC) responsible for the reduction of oxygen to water [25].

The genome of the anammox bacterium *Ca. B. pituitae* encodes a gene homologous to *kuste2479*, *bpit_14680*. The product of this gene, hereon called BpMHAO, has an unknown function and the specific roles of its individual domains also remain unexplored. To address this, our study aims to elucidate the functional contributions of each domain by producing truncated forms of the protein and characterizing them biochemically, enzymatically and spectroscopically truncated forms of the protein.

## Results and Discussion

### Amino acids sequence analysis and model prediction

The genome of *Ca. Brocadia pituitae* contains twelve paralog gene locus, which code for proteins homologous to the hydroxylamine oxidoreductase family. The *bpit_14680* gene codes for an 840 amino acids protein, BBO17176 (NCBI code), hereafter named BpMHAO. The first 25 amino acids of its sequence were predicted as a signal peptide using SignalP6.0 [26]. The presence of signal peptides is a common feature found not only in other HAO-like proteins from *Ca. B. pituitae* but also in homologous proteins from other anammox bacteria, such as *Ca. K. stuttgartiensis* [27]. Although HAO-like proteins were predicted to be located inside the anammoxosome [28] it remains challenging to definitively link the presence of signal peptides in these *Ca. B. pituitae* proteins, as well as in the other anammox bacteria, with their specific localization within the cell, such as the periplasmic space or the anammoxosome.

Following the signal peptide, we found a unique combination of two domains, a multicopper domain, MCo-like, fused to the HAO-like domain (Figure 1A).

**Figure 1.**
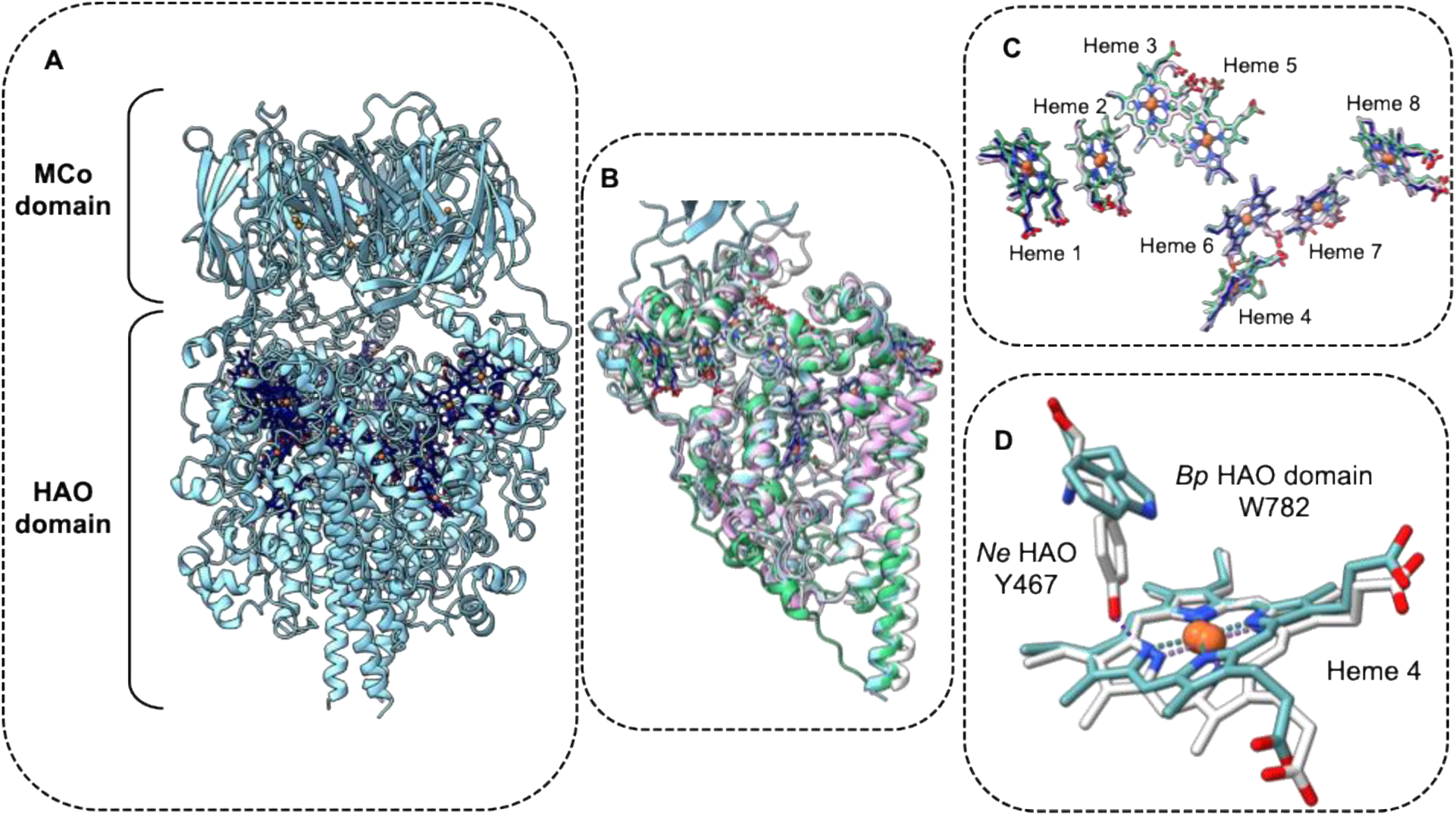
AlphaFold3 prediction of the structure of the BpMHAO protein with the copper and heme co-factors. A) Structure prediction of a trimer of the MHAO from Ca. B. pituitae B) HAO-like domain from BpMHAO superimposed with the HAO from N. europaea (PDB: 1FGJ, white), Ca. B. fulgida (PDB 6T5E, pink) and Ca. K. stuttgartiensis (PDB 6H5L, green). C) Heme distribution in MHAO in trimeric organization (blue) superimposed with the hemes of HAO from N. europaea (white). D) Structure of heme 4, the catalytic heme, superimposed with the heme 4 HAO from N. europaea (PDB: 1FGJ), highlighting the covalently bound tyrosine to the heme porphyrin. The structural analysis and the images were prepared with ChimeraX [29,30].

A BLAST [31] search using the amino acid sequence of BpMHAO showed that, as of July 2026, proteins with this combination of domains was exclusive to organisms belonging to the Planctomycetota phylum, with homologous genes present in all anammox bacteria with sequenced genomes, except for the *Ca. Scalindua* genus.

The amino acid sequence analysis showed that the MCo-like domain of BpMHAO (residues 27 to 305) has high homology with the members of the small laccase family, such as the one from *Streptomyces viridosporus*, with a 32% identity (Figure 2A). The copper binding motifs that are characteristic of the members of this two-domain multicopper bacterial laccases family, **H**_98_x**H**_100_, **H**_143_x**H**_145_, <u>H</u>_238_xx**H**_241_x**H**_243,_ and **H**_285_<u>C</u>_286_**H**_287_xxx<u>H</u>_291_ (T1 copper binding histidines are underlined whereas the trinuclear center binding histidines are in bold, BpMHAO numbering), were fully conserved in the amino acids sequence [32] (Figure 2A). When compared to the same domain of homologous proteins from anammox bacteria, we observed an increase in the amino acid sequence identities, ranging from 67 to 80%. The highest identity was found with the MCo-like domain of the homologous protein from *Ca. Brocadia fulgida*. Curiously, in the MCo-like domains of those proteins from the genus *Ca. Kuenenia*, the His145 and His287, that are predicted to coordinate the TNC center in *B. pituitae*, are replaced by a tyrosine and a glutamine, respectively (Figure 3). Nevertheless, there are examples in the literature, such as stellacyanin from *Cucumis sativus*, in which the copper is also coordinated by a glutamine [33].

**Figure 2.**
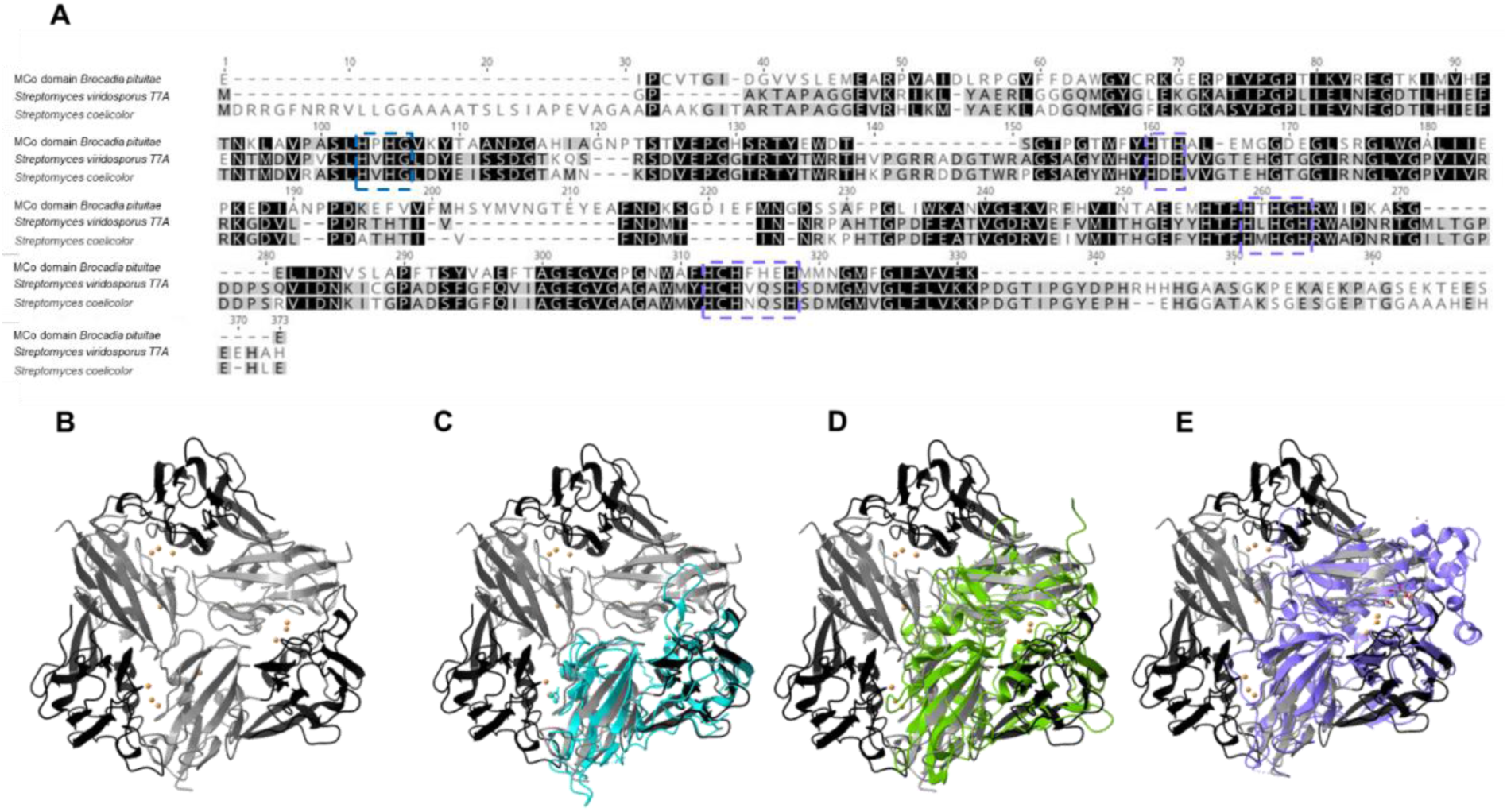
Mco-like domain of Ca. Brocadia pituitae. A) Amino acids sequence alignment of the MCo domain from Ca. B. pituitae with the small laccase from Streptomyces viridosporus and Streptomyces coelicolor. The dashed blue rectangle highlights the conserved binding motifs for the copper centers. B) AlphaFold3 prediction model structure of the MCo-like domain of Ca. B. pituitae in a trimer organization with the four copper atoms per monomer. In black is represented the N-terminal part of the proteins and in grey the C-terminal. C) AlphaFold3 prediction model structure of the MCo-like domain of Ca. B. pituitae superimposed with the 2dMCo small laccase from Streptomyces viridosporus T7A in blue (PDB:3TAS), (D) the 3dMCO CueO of E.coli (PDB: 3NSF) in green and the 6dMCO from Saccharomyces cerevisiae (PDB: 1ZPU) in purple. The structural analysis and the images were prepared with ChimeraX [29,30].

**Figure 3.**
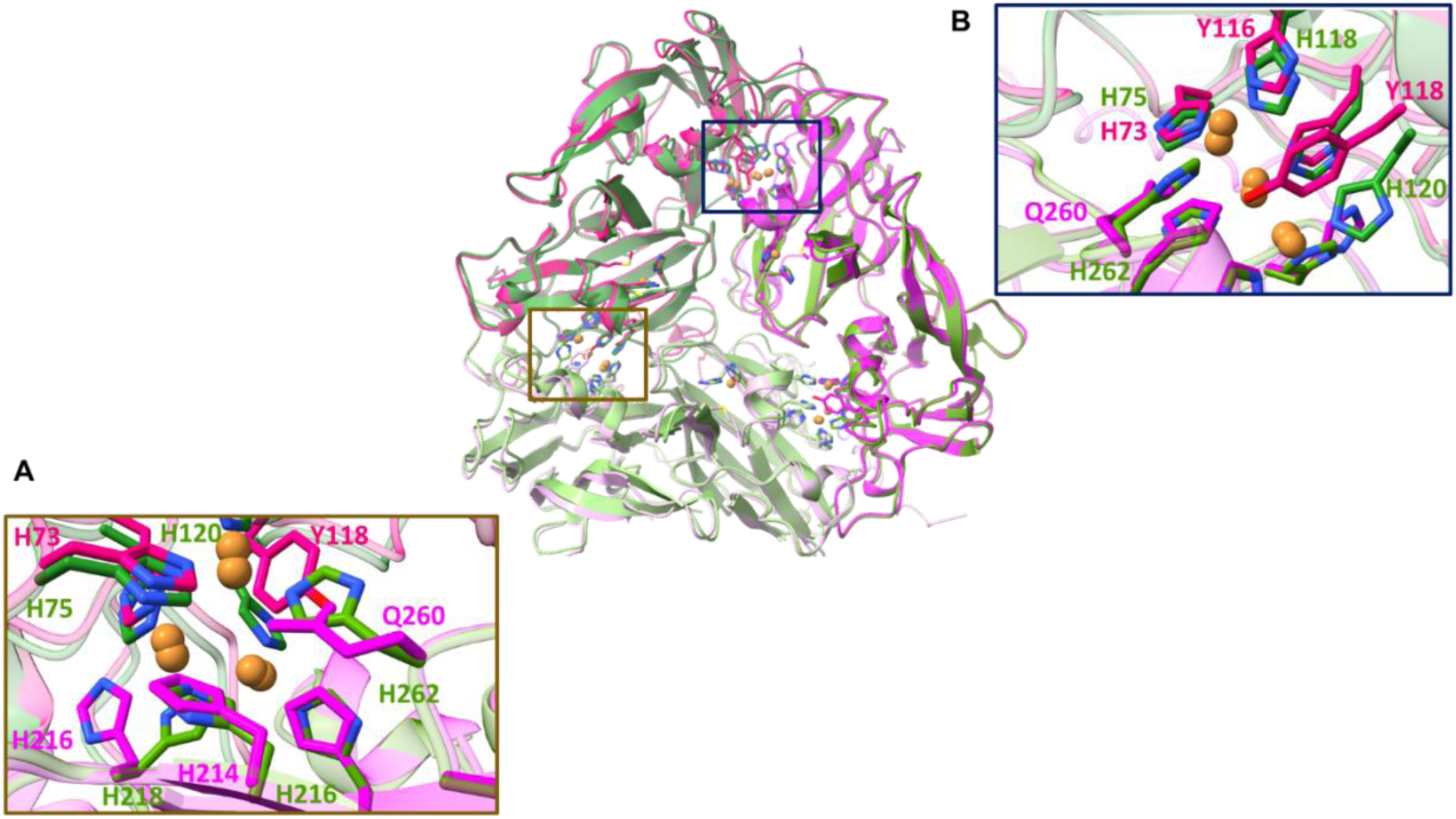
AlphaFold3 prediction of the model structure of the MCo-like domains of MHAO protein from Ca. K. stuttgartiensis (pink) and Ca. B. pituitae (green) in a trimeric organization. The different shades of pink and green correspond to different monomers for each MCo-like domain. Panels A) and B) represent the trinuclear (TNC) center with the respective copper ligands. The structural analysis and the images were prepared with ChimeraX [29,30].

AlphaFold3 [34] was used to perform the structure prediction of the *Ca. B. pituitae* MCo-like domain, assuming the presence of the predicted four copper atoms per protein monomer. This helped us to elucidate the position for each copper site as well as confirm their respective amino acids’ coordination (Figure 2B). As in other members of the small laccase family, the T1 center is inserted in a “cupredoxin-like” environment with a typical Greek key β-sandwich with parallel and antiparallel β-strands [35].

Structurally, MCo proteins can be classified into three different types, 2dMCo, 3dMCo and 6dMCo, based on the number of domains and on the location within the domain where the copper binding amino acids are located [32,36]. The members of the small laccase family usually belong to the 2dMCo family [37]. In agreement with what was observed in the amino acids sequence alignment, also the analysis of the model structure indicated that the BpMAHO MCo-like domain belongs to the 2dMCo, in particular to the type B family, which is characterized by the coordinating amino acids of the T1 copper center being in the C-terminal region of the monomer, whereas for the trinuclear center (T2 and T3), the eight coordinating amino acids of the copper ions are shared between two adjacent monomers, four from each monomer, as shown in Figure 3 [32]. As in other type B 2dMCo, the T1 and TNC sites are linked via a conserved T1-H_285_C_286_H_287_-T3 protein pathway, already observed in the amino acids sequence analysis, which should ensure an effective electron transfer between the two sites [32,38,39].

Based on this information, we built an oligomeric model of this domain, assuming a trimeric quaternary structure, characteristic of 2dMCo family of proteins. The resulting model structure was superimposable with the crystallographic structures of canonical 2dMCo-type small laccase proteins family, such as the one from *Streptomyces viridosporus* (PDB: 3TAS), with an r.m.s.d. of 5.3 Å (Figure 2C). As expected, when the model structure of the MCo-like domain was compared with the crystallographic structures of proteins members of the 3dMCO or 6dMCo types, such as the multicopper oxidase CueO from *E. coli* (PDB: 3NSF) or the Fet3p from *Saccharomyces cerevisiae* (PDB: 1ZPU), r.m.s.d. of 11.6 Å or 9.4 Å were obtained, respectively, (Figure 2D and E) [40,41].

The HAO family, on the other hand, is represented by a very broad number of enzymes that, despite having very similar three-dimensional structures, with eight *c*-type hemes, may have different catalytic functions that range from, for example, hydrazine and hydroxylamine oxidases to nitrite, nitric oxide or hydroxylamine reductases [12,42–46]. In *Ca. K. stuttgartiensis*, one of the HAO-like paralogs, Kustc1061, exhibits hydroxylamine oxidase activity, producing nitric oxide as a reaction product [11], whereas the homologous HAO from *Nitrosomonas* (*N.*) *europaea* (aerobic bacterium that oxidizes ammonium to nitrite) catalyzes the oxidation of hydroxylamine to nitrite [47].

Based on the amino acids sequence analysis of the HAO-like domain of BpMHAO (residues 325 to 840), we identified the expected binding motifs for the eight *c*-type hemes (CxxCH) that are characteristic of HAO enzymes. It shares high identity with the HAO-domains of homolog proteins from other anammox bacteria, such as *Ca. B. fulgida,* with 85% identity (Figure S1), but it is also similar (27% identity) to the HAO enzyme from *N. europeae*.

AlphaFold3 [48] was also used to build the model of the HAO-like domain, including the eight *c*-type hemes predicted by the amino acids sequence (Figure 1A). The overall model structure of BpMHAO HAO-like domain was very similar to the crystallographic structures of other HAO proteins, and it was superimposable with the homologs from *N. europaea* (PDB 1FGJ), *Ca. B. fulgida* (PDB 6T5E) and *Ca. K. stuttgartiensis* (PDB 6H5L), with r.m.s.d.s of 1.0 Å (Figure 1A), 1.0 Å and 0.7 Å, respectively. As was observed in the crystallographic structures of these HAOs, except for heme 4, the catalytic one, which is penta-coordinated with a histidine in the axial position (His499, *Ca. B. pituitae* numbering, excluding signal peptide) the remaining seven hemes are hexa-coordinated. Another important difference between the BpMHAO HAO-like domain model and the *Ca. B. fulgida* and *N. europaea* HAO structures was the absence of the tyrosine residue binding the porphyrin ring of heme 4, corresponding to Tyr449 in *Ca. B. fulgida* and Tyr467 in *N. europaea*. Structurally, this is a very interesting feature among canonical HAO enzymes because it involves a covalent cross-link between the tyrosine from one monomer and the catalytic heme of an adjacent one, which is only possible due to the quaternary arrangement of these proteins. This feature is responsible for the structural distortion of the catalytic heme, as observed, for example, in the *N. europeae’s* HAO structure (Figure 1C). Also, from the catalytic perspective, the presence of this porphyrin-binding tyrosine was proposed to be indicative of the ability of these enzymes to perform oxidation reactions [12]. Instead, in the HAO-domain of BpMHAO, this position is occupied by a tryptophan, W782 (*Ca. B. pituitae* numbering, excluding signal peptide). This substitution (tyrosine by tryptophan) is not exclusive of the HAO-domain of BpMHAO, being present in some HAO homologs from anammox bacteria with sequenced genomes (Figure S1). Contrary to tyrosine, this tryptophan is unable to bind to the porphyrin of the catalytic heme, likely preventing its distortion, and allowing its reductive character (see enzymatic activity section below).

As the crystallographic structures of both 2dMCo small laccases and HAO enzymes were trimers, we again utilized AlphaFold3 to predict and build the trimeric structure model of the full-length protein (Figure 1A). The model with the best score had an “*ice-cream-like”* cone-shaped structure formed by the HAO-like domain with the MCo-like domain on top. Surprisingly, this arrangement maintains the structures of each individual domain obtained separately. This is possibly related to the presence of an extensive 14 amino acids loop (from residue 279 to 292) loop between the two domains (Figure 1A and S2). The analysis of the per-residue confidence scores (pLDDT) provided by AlphaFold3 in that region is lower than 50, which indicates a considerable degree of uncertainty and corroborates the existence of some flexibility between the two domains. In a scenario where the MCo and HAO-like domains cooperate for the full-length enzyme’s activity, this will require inter-domain crosstalk, for example, inter-domain electron transfer. In this modeled domain organization, the shortest distance between the nearest centers of the MCo and HAO domains (T2 copper site and heme 2) is approximately 33 Å. This separation is too long to allow efficient electron transfer. Nevertheless, the flexibility of the interdomain linker could allow conformational rearrangements that transiently bring the two redox centers into closer proximity, thereby overcoming this limitation.

### Biochemical characterization

The MCo domain of BpMHAO from *B. pituitae* was successfully produced in *E. coli* and purified to homogeneity (Figure 4B). A molecular mass of ∼35 kDa, determined by SDS-PAGE, agreed with the prediction based on the amino acid sequence, which was 31 kDa. The copper content of the purified MCo-like domain was 2.92 ± 0.04 Cu/protein, which was lower than the expected 4 Cu/protein. Low copper content was also observed in other members of the MCo family [41,49]. The production of the BpMHAO HAO-like domain was attempted in both *E. coli* and *Shewanela oneidensis*, but the yields were not sufficient to proceed with the purification process. Therefore, we pursued the evaluation of the enzyme’s substrate using the soluble fraction of the cellular extracts, obtained after ultracentrifugation, hereafter named SoHAO and EcHAO for *S. oneidensis* and *E. coli* extracts, respectively. The presence of the target protein in these extracts was confirmed by western blot analysis, with an antibody against the His-Tag that was present at the C-terminal part of the protein (Figure 4D). The western blot shows more than one band, which was attributed to protein degradation over time. Therefore, it was assumed that the functional protein was the one with the correct predicted molecular mass of ∼67 kDa (accounting for the 8 *c-*type hemes). The western blot was also used to estimate the amount of protein present in the cell extract, which was determined as 0.012 mg/mL and 0.013 mg/mL, that correspond to 0.04 ± 0.02 % and 0.04 ± 0.02 % of the total amount of protein of EcHAO and SoHAO, respectively (Figure S4). Cell extracts produced with expression vectors without the gene for the BpMHAO HAO-like domain were also produced to be used as controls (named SoControl and EcControl for *S. oneidensis* and *E. coli* extracts, respectively).

**Figure 4.**
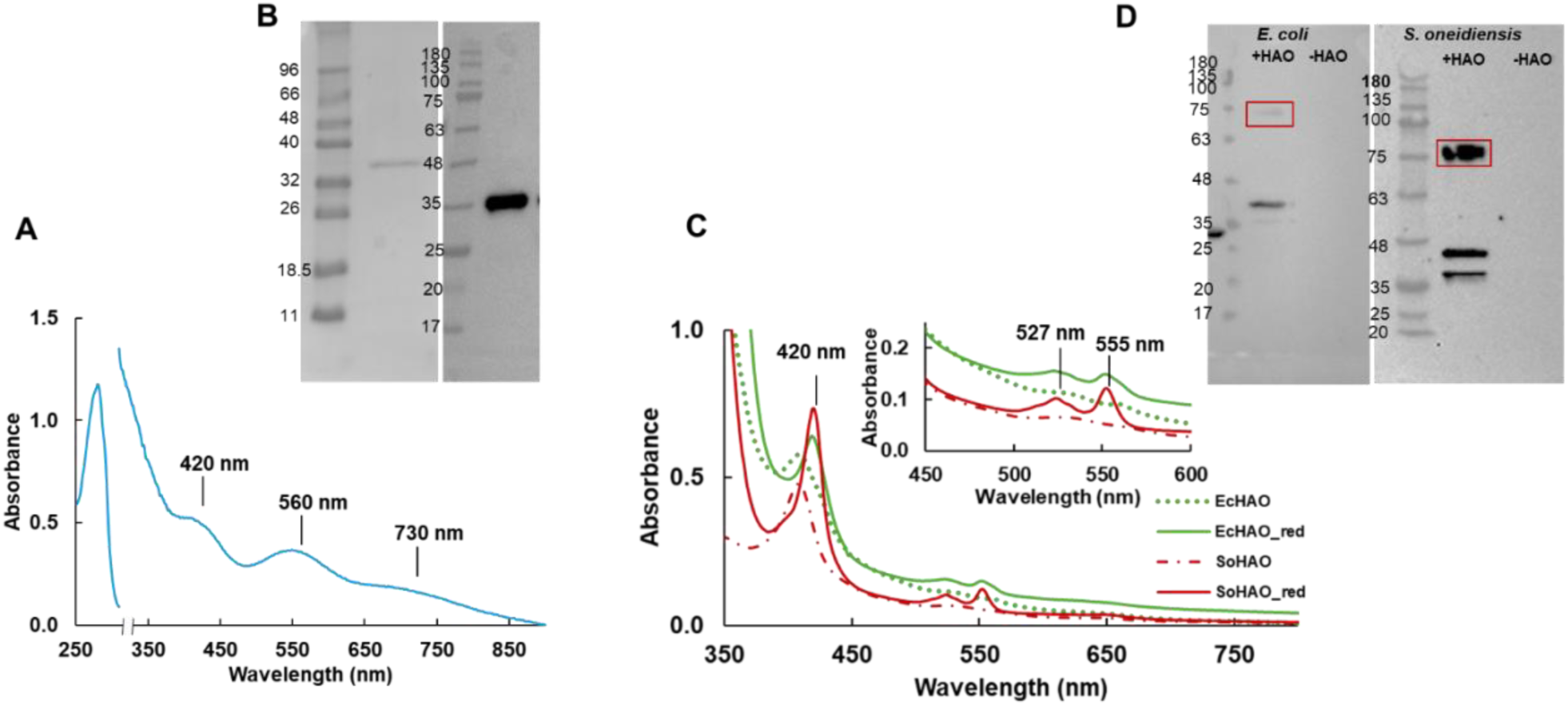
UV-visible spectroscopic features of the MCo domain, EcHAO, and SoHAO. A) UV-visible spectrum of the pure MCo-like domain in the oxidized state in 50 mM Mops pH 7.5. B) SDS–PAGE (left) and western Blot (right) of pure MCo-like domain C) UV-visible spectrum of the EcHAO (green) and SoHAO (red) in oxidized (dashed lines) and reduced form (solid lines). The spectra were acquired in 50 mM of sodium phosphate pH 7.4. The reduction of both samples was performed by adding small amounts of sodium dithionite crystals. D) Western blot of the cell extracts of E. coli and S. oneidensis cells transformed with (+HAO) and without (-HAO) the plasmid containing the gene for the HAO-like domain. The red square corresponds to the gel band considered for HAO-like domain in the active form.

### UV-visible and EPR Spectroscopies

The UV-visible spectrum of the MCo-like domain of BpMHAO in the as-purified state presented absorption bands with maxima at 428, 560 and 695 nm (Figure 4A). These spectral features are uncommon for the members of the small laccase family, whose UV-visible spectra has usually maxima at ∼600 nm, arising from the contribution of the T1 center, and at ∼330 nm, which results from the contribution of the T3 center [41,50–53]. Interestingly, a comparable UV-visible spectrum was observed in the single domain cupredoxin from *Acidithiobacillus ferrooxidans,* which was attributed to the tetragonal distorted T1.5-like center, where the band at 438 nm was attributed to the S(Cys) → Cu σ charge-transfer transition, whereas the band at 568 nm was a result of the S(Cys) → Cu π charge-transfer transition, and the broad peak around 690 nm likely corresponded to d-d transition bands [54]. Contrary to what was observed with the *Acidithiobacillus ferrooxidans* cupredoxin, an absorption band with maximum at 324 nm appeared in the UV-visible spectrum of the MCo-domain of BpMHAO when going from pH 3 to pH 7.5. After this point the UV-visible spectrum remained the same (Figure S3). In similar proteins, these absorption band is usually attributed to the contribution of the T3 type center [25].

Nevertheless, a UV-visible spectrum, with maxima at 425, 570 and 700 nm, was observed for the “oxy” form of hemocyanin from *Carcinus aestuarii*. This spectrum was attributed to the protein’s binuclear copper center coordinated by six histidines and a peroxo-bridge [25,55,56]. In oxi-hemocyanin, the absorption maximum at 570 nm was attributed to the peroxo-copper charge transfer, whereas the maximum at 425 nm was related to the histidine-to-copper charge transfer [56].

The EPR spectrum of BpMHAO MCo-like domain (Figure 5A) displayed a signal pattern typical of the presence of monomeric copper centers with the characteristic hyperfine splitting resonances arising from the copper nuclear spin, I=3/2. By spectral simulation, we obtained two sets of g values and hyperfine constants, *g* = 2.045, 2.045, 2.206 and A_//_ = 100 x 10^-4^ cm ^-1^ and *g* = 2.071, 2.071 and 2.257 and A_//_ = 194 x 10 ^-4^ cm ^-1^, consistent with the presence of T1 and T2 type centers, respectively. These parameters are similar to those reported for other multicopper proteins, namely from the laccase family [57] The absence of an EPR signal attributable to the T3 center agreed with an antiferromagnetic coupling of the two copper ions, expected for this type of copper centers and usually related to the presence of a bridging hydroxyl ligand [56]. A species tentatively assigned to the T3 center was observed during the redox titration (see below).

**Figure 5.**
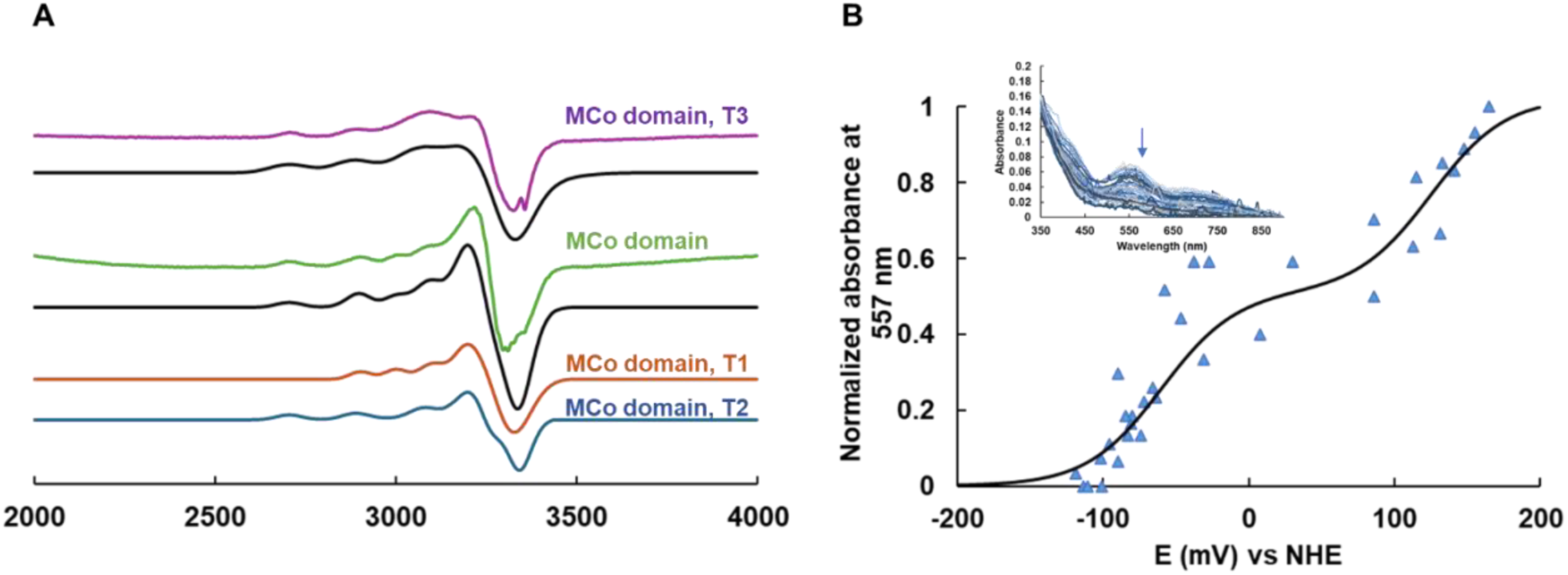
EPR spectra and redox titration followed by UV-vis. A) EPR spectra of 100 µM of MCo-like domain in the as-purified state (green spectrum) and the sample prepared at 7mV, collected during the redox titration (purple), in 50 mM of MOPS pH 7.5, as prep (green). The black spectra correspond to the simulations of each spectrum. In the case of the as purified sample, the contribution of T1 and T2 centers used for the simulation is represented in orange and blue lines. Microwave Frequency 9.4 GHz, temperature 15 K, Modulation amplitude 10G, power 2 mW. B) Anaerobic redox titration followed by UV-visible spectroscopy at 557 nm of the MCo-like domain from BpMHAO (blue triangles) in 50 mM MOPS pH 7.5. The solid line represents the Nernst equation model for two consecutive monoelectronic steps as described in the Materials and Methods section. The inset represents the UV-visible spectra acquired from two experiments.

Therefore, despite the fact that we cannot discard the contribution of the T1 center to the UV-visible spectrum of the MCo-like domain of BpMHAO, namely because our EPR data confirm that both T1 and T2 centers are present in the “as-prepared” oxidized protein, by combining the results obtained from both UV-visible and EPR monitored titrations (see below), we propose that the UV-visible spectrum is mainly dominated by the contribution of the T3 copper center, possibly with a peroxo-bridge, in a configuration similar to the “peroxy intermediate” described by *Solomon et. al.* [58]. The presence of this peroxo-bridge between the two copper ions is also consistent with the antiferromagnetic coupling of center T3 and, consequently with the absence of EPR signal of this species in the as purified sample. Furthermore, the incubation of the “as purified” sample with hydrogen peroxide did not produce any modification of the UV-visible spectrum, which excludes the lack of contribution of any center, namely T1, to the spectrum of the purified protein. As for similar proteins, the T2-type center does not contribute to the UV-visible spectrum[25].

The UV-visible spectra were obtained for the EcHAO and SoHAO cell extracts, displaying typical features of *c*-type heme containing proteins, with maxima at 420, 527 and 550 nm in the reduced state, corresponding to the Soret, β and α bands, respectively (Figure 5C).

### Reduction potential of MCo-like domain

The reduction potential of the BpMHAO MCo-like domain was obtained by anaerobic titration monitored by UV-Visible and EPR spectroscopies (Figure 5A, B). The initial sample of the EPR titration, prepared at 136 mV, showed no signal, indicating that, at this potential, both T1 and T2 centers were already reduced. This result is not unexpected as T1 and T2 centers of other MCo proteins usually have reduction potentials much higher than 136mV, e.g., the reduction potential of T1 centers can range between ∼ 300 mV to 800 mV [25,59,60]. The only EPR-active species observed during this titration was obtained in the sample prepared at 7 mV (Figure 5A). The spectrum had features similar to a mononuclear copper site, and by spectral simulation we obtained g values of 2.073, 2.073, 2.270 and A// = 185 x 10^-4^ cm^-1^, which were different from those of T1 and T2, and, therefore, we attributed them to a mixed-valence T3 center, with a delocalized charge to one of the coppers, Cu^2+^Cu^1+^, detectable until 60K (data not shown).

The UV-visible titration was monitored at 557 nm (Figure 5B). Taking into consideration that both EPR and UV-visible monitored titrations were performed in a similar potential range, and that in the titration monitored by EPR, the T1 and T2 centers were not observed, we considered, as proposed in the previous topic, that the main contributor to the UV-visible spectrum of the MCo-like domain of BpMHAO was the T3 center. Thus, a Nernst equation for two consecutive monoelectronic transitions (Cu^2+^Cu^2+^ → Cu^2+^Cu^1+^ → Cu^1+^Cu^1+^) was used to calculate the reduction potential of the binuclear copper center, resulting in 125 mV ± 10 mV and −60 ± 10 mV. The reduction potential of T3 copper centers can also vary but are usually higher than the ones calculated for the T3 center from MCo-like domain from the BpMHAO protein [61].

### Enzymatic Activities

#### MCo-like domain

Different functions have been assigned to multicopper oxidases in the literature (17). Because the MCo-like domain of BpMHAO has features related to the small laccase family, we tested different substrates that are usual for this family of enzymes. Different pH values and temperatures were also evaluated to achieve the best conditions for the activity assays. Firstly, 2,2’-Azino-bis(3-ethylbenzothiazoline-6-sulfonic acid) (ABTS) and catechol were used as substrates to evaluate for laccase activity. The results showed that the reaction using catechol as substrate was very limited. Using ABTS as substrate, we calculated a reaction rate of 0.06 U/mg at pH3, while it was negligible at pH 7 (Figure S4A). These results showed that, similarly to other laccases [62,63], the optimum pH for the MCo-like domain was pH 3. Nevertheless, a protein precipitate started to form after a few seconds, which did not occur with the control assays performed with the fungal laccase from *Tramates versicolor* (Figure S4B). These results showed that, despite its ability to oxidize ABTS, the MCo-like domain was not stable at the optimal conditions required for the enzymatic reaction.

Despite the low homology between the MCo-like domain of BpMHAO and the copper-containing nitrite reductases, we evaluated its ability to reduce nitrite to nitric oxide, which is indeed an intermediate in the anammox reaction, and could indicate a possible role of this domain or the full-length protein. Nitrite reductase activity assay followed that described for cytochrome *cd*_1_ nitrite reductase[64], using methyl viologen as an electron donor. Nevertheless, no enzymatic activity was observed (Figure S4C).

Finally, because the MCo superfamily includes enzymes with ferroxidase activity, such as the human ceruloplasmin, the Fet3p from *Saccharomyces cerevisiae,* and the HaLF from *Heterobasidion annosum s.l.* [63,65,66], we also evaluated the ability of the MCo-like domain to perform the oxidation of Fe^2+^ to Fe^3+^ using oxygen as a co-substrate.

Ferroxidase activity was first assessed by UV-visible spectroscopy, by monitoring the oxidation of Fe^2+^ to Fe^3+^ at 315 nm [67,68]. We observed that the MCo-like domain was able to oxidize Fe^2+^ at a rate of 0.24 ± 0.05 U.mg^-1^. These reaction rates are in the range with of those obtained for metallooxidases [39]. By titrating the enzyme with consecutive additions of FeCl_2_, we observed that the MCo-like domain was able to oxidize up to 40 Fe^2+^/protein until we observe insoluble iron particles in the reaction mixture (Figure 6A). In a control assay performed in the absence of the enzyme, iron precipitation was observed upon the initial addition of iron (Figure 6B). An additional control was also performed in the absence of oxygen, which showed that the iron oxidation does not occur (Figure 6C).

**Figure 6.**
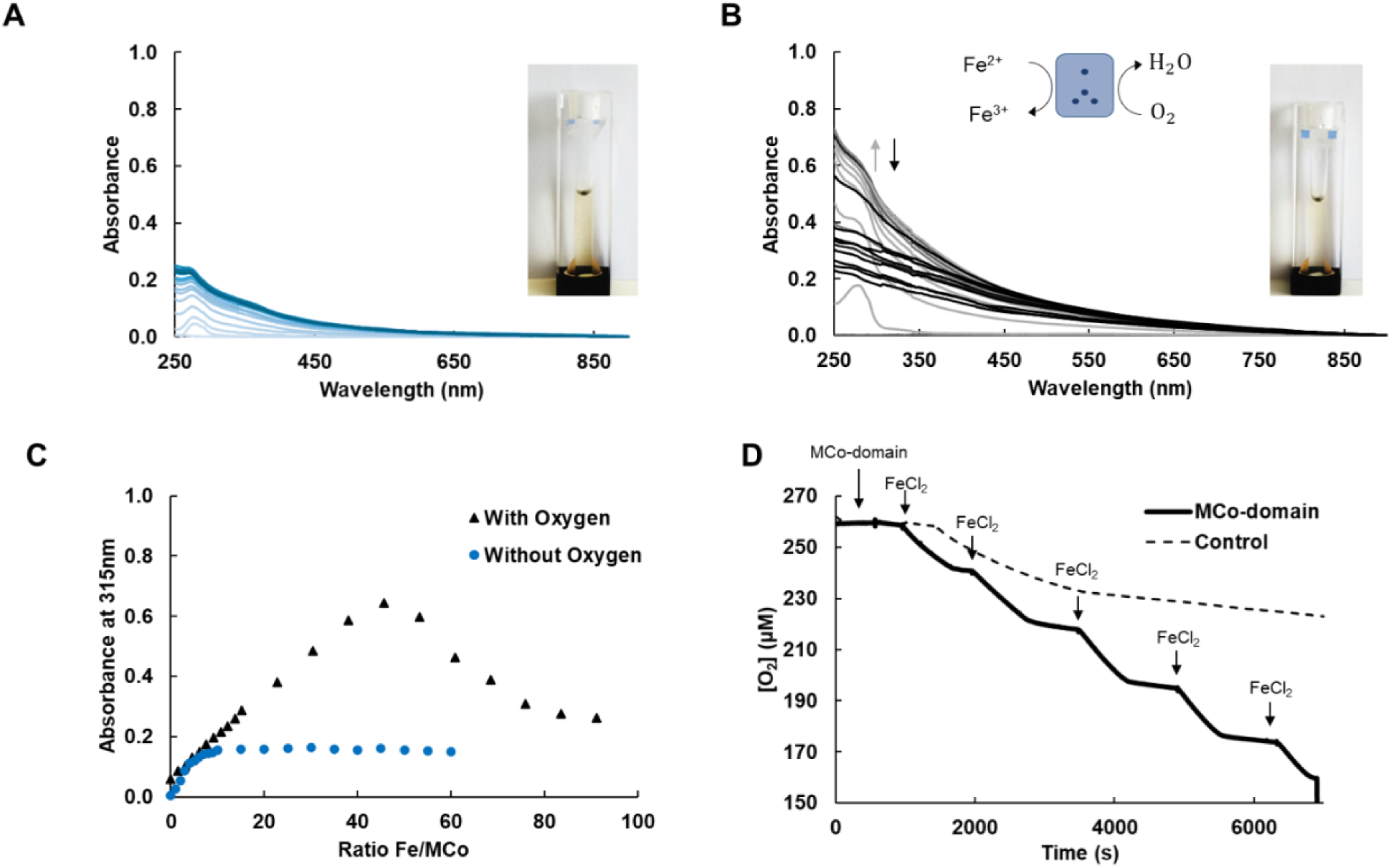
Ferroxidase activity of the MCo-like domain. A) Control assay performed by titrating FeCl_2_ to a 50 mM of MOPS pH 7.5 with 150 mM of NaCl monitored by UV-visible spectra. B) Titration of FeCl_2_ (up to 300 µM) to a 50 mM of MOPS pH 7.5 with 150 mM of NaCl containing 5 µM of MCo-like domain, monitored by UV-visible spectra. C) Representation of the absorbance at 310 nm of the same experiment as in (B) in the presence of oxygen (blue dots) and without oxygen (black triangles). D) Oxygen consumption assay performed as in (B) in the presence of 5 µM of MCo-like domain (black line) and without the protein (black dashed line). The (A), (B) and (C) are representative data of 3 replicates whereas the (D) represents one of four replicates.

As oxygen should act as a co-substrate for this reaction, we performed a similar assay but monitoring it using a Clark-type electrode, aiming not only to monitor the O_2_ consumption rate during iron oxidation but also to calculate the iron/oxygen stoichiometry. In this case, stepwise additions of 50 µM of FeCl_2_ to the reaction mixture resulted in the consumption of ≈ 20 µM of oxygen (Figure 6D), pointing to a stoichiometry of 2:1 between iron and oxygen, and indicating an incomplete reduction of oxygen to hydrogen peroxide and not to water, as observed in other multicopper enzymes [39].

The ferroxidase activity of Fet3p was attributed to three key acidic residues: D283, D409 and E185 (Fet3p numbering) [63]. Negatively charged amino acids oriented towards the T1 center, were suggested to be directly involved in the oxidation of Fe^2+^ to Fe^3+^. In HaLF protein, substitution mutants (A227E, F344D, R422Y and D475A – HaLF numbering) of the corresponding residues confer ferroxidase activity, which is absent in the wild-type protein [23,69]. A close inspection of the superposition of the MCo-like domain with Fet3p shows that the possible amino acids that could interact with the iron could be E167, E169 and E211 (Figure S5). This arises from being the only negatively charged amino acids close to the T1 copper center, which could be involved in the interaction with iron.

#### HAO-like domain

Similar to the MCO family, the HAO family also has a very wide range of enzymatic functions associated with it and, consequently, a broad range of potential substrates. However, when focusing on the hypothetical role of the full-length enzyme in *Ca. B. pituitae*’s anammox process, we tested both its putative nitrite reductase and hydroxylamine oxidase activities (putative NO producer). The nitrite reductase activity was similar to that described for one of the HAO-like paralogs from *Ca. K. stuttgartiensis* (Kustec0458/7) [12], and also similar to that performed using the MCo-like domain. These assays, using methyl viologen as an electron donor, led to a rate of 0.4 ± 0.2 µM of methyl viologen. s^-1^ for both EcHAo and SoHAO. These values were comparable to the corresponding control experiment with EcControl, 0.3 ± 0.03 µM of methyl viologen. s^-1^, and SoControl, 0.5 ± 0.1 µM of methyl viologen. s^-1^ (Figure S6A). Therefore, this HAO-like domain is unable to perform the reduction of nitrite. The oxidation of hydroxylamine was also evaluated, and in this case, we used horse heart cytochrome *c* as an electron acceptor, as was shown before [70]. We obtained rates of 0.02 ± 0.01 and 0.13 ± 0.03 mM of reduced Cyt*c.*s^-1^, for EcHAO and SoHAO, respectively. Again, the rates observed for the control assays with EcControl and SoControl were similar, indicating that this domain could not perform oxidation of hydroxylamine (Figure S6B).

Finally, we tested for hydroxylamine reductase activity for both cell extracts containing the HAO-like domain. The assays were performed using methyl viologen as electron donor, and the results showed that both EcHAO and SoHAO cell extracts were able to perform hydroxylamine reductase activity at a rate of 240 ± 25 µM of methyl viologen. s^-1^.mg^-1^ and 660 ± 115 µM of methyl viologen. s^-1^.mg^-1^, for EcHAO and SoHAO, respectively (Figure 7A), considering the amount of HAO-like domain calculated from the western blot (see materials and methods). EcControl and SoControl also presented residual hydroxylamine activity, possibly due to the presence of enzymes with this capability in both cell extracts. To confirm that the hydroxylamine reduction reaction was due to the presence of the BpMHAO HAO-like domain in the cell extracts and not an artifact, we performed several assays, similar to the previous ones, but with increasing amounts of cell extract (0.1 mg/mL to 0.5 mg/mL of total protein content). The results showed a linear increase in the hydroxylamine reduction rate with the amount of cell extract (HAO-like domain concentration), as expected for an enzymatic driven reaction (Figure 7B). Although hydroxylamine reductase activity is not common in HAO-like proteins, some can perform this reaction as a non-physiological one, but with lower rates [12]. The higher rates observed for the truncated HAO-like domain point to its physiological role. To calculate the Michaelis-Menten parameters for this enzyme, we performed similar assays but using a range of hydroxylamine concentrations up to 5 mM of hydroxylamine (Figure 7C).

**Figure 7.**
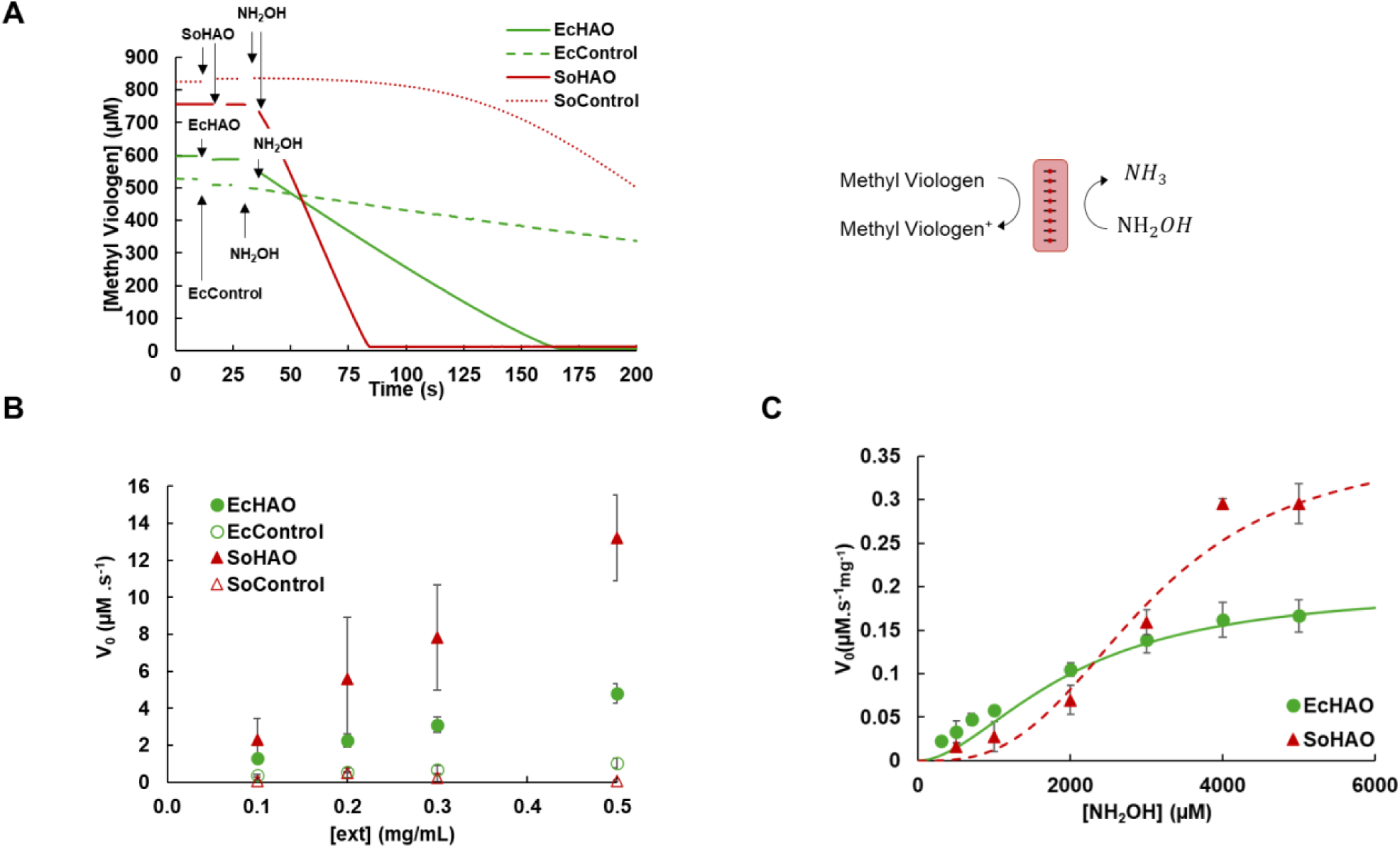
Hydroxylamine reductase activity of the E. coli and S. oneidensis cell extracts. A) Hydroxylamine reductase activity of 0.5 mg/mL of EcHAO (green line), EcControl (green dashed), SoHAO (red line) and SoControl (red dotted) with 1mM of hydroxylamine. Hydroxylamine reduction was monitored indirectly via the oxidation of methyl viologen at 730 nm (ε_730nm_ = 2.13 mM^-1^.cm^-1^) in anaerobiosis, in 50 mM Tris-HCl pH 7.5. B) Representation of the enzymatic activity (µM of methyl viologen. s^-1^) as a function of cell extracts concentration (EcHAO, solid green dots), EcControl) (open green dots), SoHAO (red triangles) and SoControl (open red triangles). C) Michaelis-Menten representation of the enzymatic activity (µM of methyl viologen. s^-1^) as a function of hydroxylamine concentration. The data for EcHAO and SoHAO are represented in green and red, respectively. The dots represent the experimental data points and solid lines represent the theoretical Hill curve adjusted to the experimental data. The experimental points are representative of three replicates.

A sigmoidal behavior was observed for both cell extracts, which indicates that the reaction follows a cooperative mechanism, and therefore, we utilized a modified Michalis-Menten equation in order to calculate the Hill coefficient. For EcHAO we obtained a *K_m_* of 2.0 ± 0.2 mM, a *V_max_* of 18 ± 7.0 µM s^-1^ and a Hill’s coefficient of 1.4, which indicates a moderate cooperativity, whereas, for the SoHAO we obtained a *K_m_* of 3.0 ± 0.2 mM, a *V_max_* of 28 ± 7.0 µM s^-1^ and a Hill’s coefficient of 2.8, which indicates a highly cooperative mechanism. These values indicate that the HAO-domain has a higher affinity to hydroxylamine when compared, for example, with Hybrid Cluster Protein from *E. coli,* which has hydroxylamine reductase activity, *K_m_* of 38.9 ± 3.5 mM [71].

During these assays, we failed to detect the production of ammonium from the reduction of hydroxylamine using colorimetric commercial kits. This could be attributed to other enzymes present in the cell extracts that are able to consume ammonium.

## Conclusion

In this study we present the BpMHAO protein from *Ca. B. pituitae.* So far, a unique member among hydroxylamine oxidoreductases due to its N-terminal multicopper domain. Here, the biochemical and spectroscopic characterization of the truncated domains (MCo-like and HAO-like) of this very intriguing protein was first reported, and consequent extrapolation of the enzymatic activity and role in the anammox reaction as a full-length protein.

Amino acids sequence alignments revealed that the MCo-like domain shares high homology with small laccase multicopper oxidases, with conserved T1, T2, and T3 copper-binding motifs. Structural modeling indicates that this domain is classified as a type B 2dMCo. Spectroscopically, the isolated MCo-like domain has atypical features for this class, displaying a distinct purple color and UV-visible spectrum with absorbance maxima at 428 nm and 560 nm, with a shoulder near 695 nm, which we attributed to a peroxo-bridged T3 copper site. Furthermore, EPR spectroscopy confirms the presence of the T1 and T2 copper sites in the as-prepared sample, and the T3 copper site in a partially reduced sample, thereby verifying the full complement of copper centers predicted for this domain. Enzymatically, the truncated MCo-like domain exhibited limited laccase activity towards ABTS and catechol but was competent in catalyzing the oxygen-dependent Fe^2+^ oxidation to Fe^3+^, oxidizing up to 40 iron atoms per protein molecule.

The soluble cell extracts, EcHAO and SoHAO, harboring the recombinant truncated HAO-like domain was confirmed and quantified by Western blot. The UV-visible spectra of both cell extracts showed the typical features arising from the presence of *c*-type hemes, with the characteristic Soret (420 nm), β (527 nm), and α (550 nm) bands in the reduced state. Enzymatic profiling of this domain in cell extracts revealed robust hydroxylamine reductase activity using methyl viologen as the electron donor. The kinetic parameters indicate atypical Michaelis-Menten behavior towards this substrate. Most importantly, the assays showed a 2:1 stoichiometry between methyl viologen and hydroxylamine, indicating that the final product of the reaction is hydrazine.

Collectively, these data suggests that the full-length BpMHAO protein functions predominantly as a hydroxylamine reductase. While the MCo-like domain possesses detectable ferroxidase activity under aerobic conditions, this activity is most probably not physiological in the context of the anaerobic lifestyle of anammox bacteria [6,72].

The BpMHAO protein likely represents a key adaptive mechanism for anammox organisms, enabling the efficient reduction of hydroxylamine, either produced as bi-product during hydrazine synthesis, or present in the outside media, and potentially facilitating its integration into the anammox process directly as hydrazine, as suggested in Figure 8.

**Figure 8.**
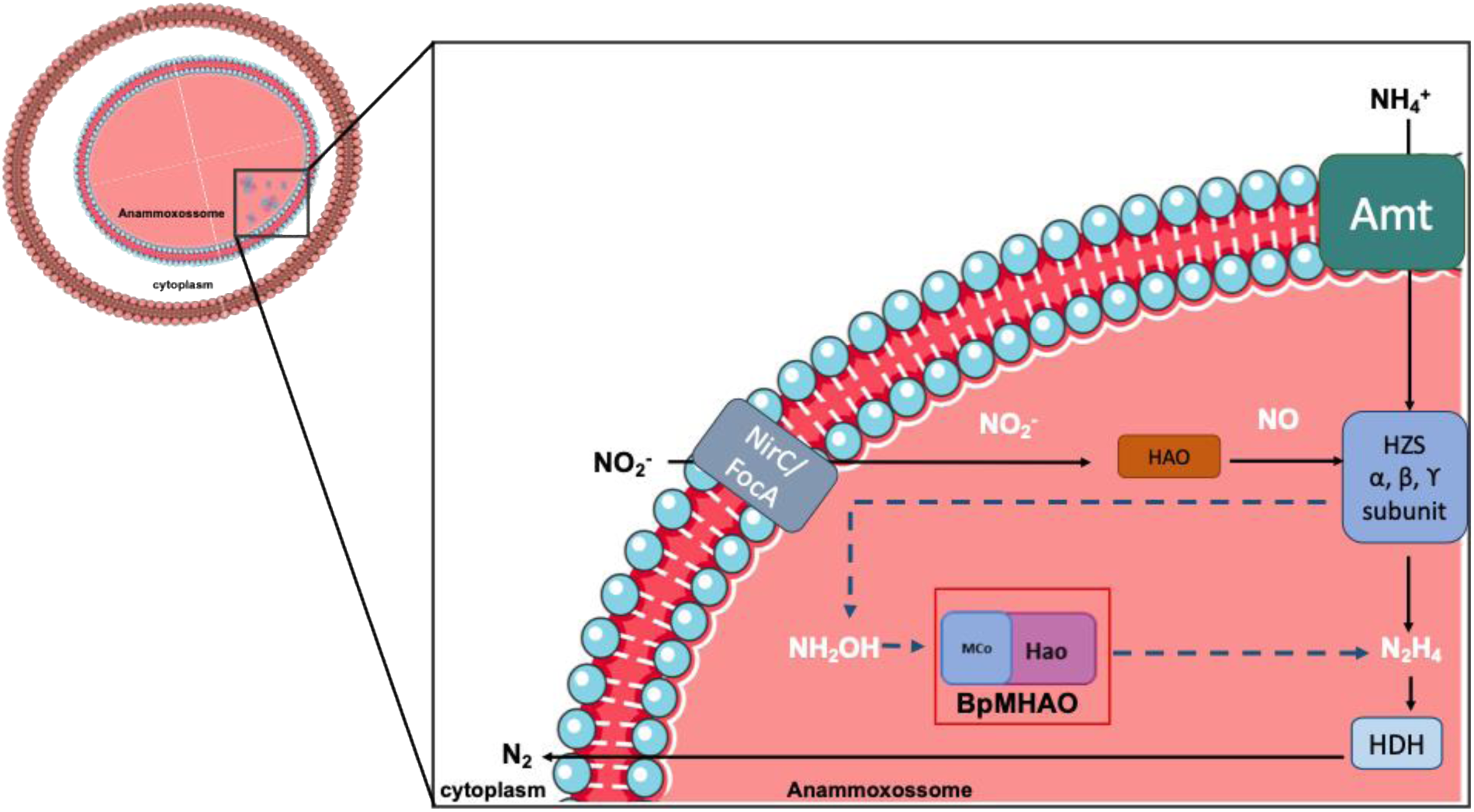
Schematic representation of an anammox bacteria cell, illustrating the proteins relevant for the anammox cycle. HAO (kustc0458 in K. stuttgartiensis) HZS-Hydrazine synthase (α, β and γ subunits, encoded by kuste2861, kuste2860, and kuste2859 in K. stuttgartiensis, respectively), HDH-Hydrazine dehydrogenase (encoded by kustc0694 and kustd1340 in K. stuttgartiensis) Transporters for nitrite (NirC and FocA) and ammonium (Amt) were already identified in this organism.

## Materials and methods

### Amino acid sequence analysis and protein structure prediction

The amino acid sequence of the full-length protein, BpMHAO (gene name *bpit_14680*), from *Ca. B. pituitae* was obtained from the NCBI database. The amino acid sequence alignments were performed using Geneious tool. The model structure predictions of both MCo-like and HAO-like domains as well as of the full-length protein, were performed using AlphaFold3 [34]. Structural analysis, visualization and image preparation were performed using UCSF ChimeraX 1.10 [29,30].

### Protein production and purification

The genes coding for the truncated multicopper (MCo-like, residues 26 to 305) and hydroxylamine oxidoreductase (HAO-like, residues 325 to 840) domains were synthesized and cloned into pET-28a(+) plasmid by GeneCust – Custom Services for Research, with codon optimization for expression in *E. coli*. A gene coding for OmpA (Outer membrane protein A) signal peptide from *E. coli* (Table 1) was added to the expression plasmid before the HAO-like domain gene to ensure the protein translocation to the periplasmic space, where the *c*-type hemes are assembled in *E. coli*. The gene constructs of both truncated domains have an histidine-tag at the C-terminal.

**Table 1.** Amino acids sequence of the signal peptides used in this study, including the one part of the original BpMHAO protein sequence.

| Signal peptide protein name | Amino Acid Sequence | Reference |
| --- | --- | --- |
| *E. coli* OmpA | MLLYAIAVLAGFAYVAQA | [74] |
| *S. oneidensis* STC | VSKKLLSVLFGASLAALALSPTAFA | [75] |
| *Ca. B. pituitae* MHAO | MFKQSRIYLP MVIVCLLFILKTGYA | This paper |

**Table 2.**
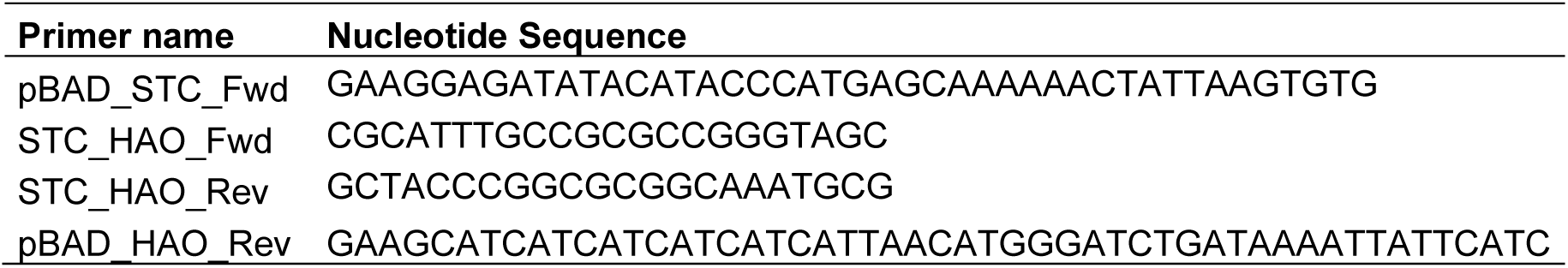
Nucleotide sequence of the primers used for the fusion of the STC signal peptide with the HAO-like domain gene.

The plasmid containing the BpMHAO MCo-like domain gene was transformed into *E. coli* BL21 (DE3) competent cells and grown in Luria Bertani (LB)-Agar plates containing 50 µg/mL of kanamycin for approximately 16h. A single colony was selected to inoculate 250 mL of LB medium supplemented with 50 µg/mL of kanamycin. The culture was allowed to grow overnight at 150 rpm and 37 °C. The overnight cultures were used to inoculate (3 % inoculum) 1 L of M9 minimal medium in 2 L Erlenmeyer flasks containing 50 µg/mL of kanamycin.

The cell cultures grew at 150 rpm and 37 °C until reaching an optical density of 0.6 at 600 nm. Then, 100 µM of isopropyl-β-D-1-thiogalactopyranoside (IPTG) and 0.1 mM of CuSO_4_ were added. The cells were grown for additional 20 h at 130 rpm and 30 °C. Cells were then harvested by centrifugation, for 10 min at 6500 x *g*, resuspended in a 20 mM Tris–HCl buffer pH 7.5 (Buffer A) and disrupted by three cycles in a high-pressure homogenizer, EmulsiFlex^TM^ – C5 (Avestin, Inc) in the presence of DNAse (Applichem). The crude extract was clarified by centrifugation at 25,000 x *g* for 20 minutes and ultracentrifugation at 138,000 x *g* for 1 hour and 30 minutes at 4 °C, to remove cell debris and membrane fractions, respectively. The supernatant from ultracentrifugation was loaded into a prepacked High-Affinity Ni-Charge Resin FF column (GenScript) pre-equilibrated with buffer A. Protein elution was performed using a gradient from buffer A to 50 mM Tris-HCl pH 7.5, 500 mM imidazole (buffer B). After this point, protein samples were analyzed by UV-visible and SDS-PAGE. Pure samples were pooled, concentrated and buffer exchanged into 50 mM Tris-HCl pH 7.5.

The production of BpMHAO HAO-like truncated domain was performed using *E. coli* C41 (DE3) competent cells, co-transformed with both the plasmid containing the gene for this domain and pEC86 containing the *c*-type hemes maturation genes *ccmABCDEFGH* from *E. coli* [73], grown in LB-Agar plates containing 50 µg/mL of kanamycin and 34 µg/mL of chloramphenicol for approximately 16h. A single colony was selected to inoculate 250 mL of LB medium supplemented with 50 µg/mL of kanamycin and 34 µg/mL of chloramphenicol. The culture was allowed to grow overnight at 150 rpm and 37 °C. The overnight cultures were used to inoculate (3 % inoculum) 1 L of 2xYT (2x Yeast extract Tryptone) medium in 2 L Erlenmeyer flasks containing 50 µg/mL of kanamycin and 34 µg/mL of chloramphenicol. Cell cultures grew at 150 rpm and 37 °C until reaching an optical density of 0.4 at 600 nm. Protein production was initiated by the addition of 100 µM of IPTG, and the media was supplemented with 0.1 mM FeCl_2_ and 1 mM 5-Aminolevulinic Acid. The cells were grown for additional 20 h at 130 and 18 °C rpm. After this point, cells were harvested by centrifugation, for 10 minutes at 6500 x *g*, resuspended in 50 mM of sodium phosphate buffer at pH 7.4 and disrupted by a single cycle in a high-pressure homogenizer, EmulsiFlex^TM^ – C5 (Avestin, Inc) in the presence of DNAse (Applichem). The crude extract was clarified by centrifugation at 25,000 x *g* for 20 minutes and ultracentrifugation at 138,000 x *g* for 1 hour and 30 minutes at 4 °C, to remove cell debris and membrane fractions, respectively.

A similar procedure was performed to prepare a parallel *E. coli* cell extract without the BpMHAO HAO-domain to be used as a control, but in this case, a pET28a(+) plasmid was used to transform the *E. coli* cells. The *E. coli* soluble fractions with and without HAO-domain were named EcHAO and EcControl, respectively.

The heterologous production of the BpMHAO HAO-like domain was also performed using *Shewanella oneidensis ΔSTC* ΔFccA (STC - Small TetrahemeCytochrome *c;* FccA - periplasmic fumarate reductase) as host, to avoid the presence of some of the most abundant periplasmatic cytochromes, STC and FccA, which should facilitate the production of a clearer cell extract. As for the plasmid used in *E. coli*, also in this case the HAO-like domain gene was fused with a signal peptide, the one from STC, by fusion PCR (the primers are described in supplementary Table 2), to promote the translocation of the HAO-like domain to the periplasmatic space. The insertion of the two genes into a pBAD plasmid was performed by NEBuilder HiFi DNA assembly Master mix (New England Bio Lab). The pBAD with the desired gene was amplified in *E. coli* DH5α (NZYtech) and purified using the NZYMiniprep kit (NZYtech).

The pBAD plasmid containing the HAO-like domain gene was transformed in *Shewanella oneidensis ΔSTC* ΔFccA by electroporation and the cells grown in LB-agar plates supplemented with 50 µg/mL of kanamycin. A single colony was selected to inoculate 250 mL of LB medium supplemented with 50 µg/mL of kanamycin and left to grow overnight at 150 rpm and 30 °C. One percent of the overnight culture was used to inoculate 2 L of Terrific Broth (TB) medium in 5 L Erlenmeyers flasks supplemented with 50 µg/mL of kanamycin. The cells were allowed to grow at 150 rpm and 30 °C until reaching an optical density of 1.3 at 600 nm. At this point, protein production was initiated by adding 0.5 mM of arabinose and the media supplemented with 1 mM 5-Aminolevulinic Acid and 0.1 mM of FeCl_2_ and the cells were grown for additional 62 h at 130 rpm and 14° C. After this, cells were harvested by centrifugation, for 10 minutes at 6500 x *g*, resuspended in 50 mM Tris-HCl at pH 7.5 and disrupted as stated above for the *E. coli* cell extracts. A similar procedure was performed but transforming *S. oneidensis* with a pBAD plasmid to serve as a control. The *S. oneidensis* soluble fractions with and without HAO-like domain were named SoHAO and SoControl, respectively.

### Protein and metal quantifications

The protein content in both isolated protein and cell extracts was obtained using the Bradford method (Bio-Rad) and bovine serum albumin (Thermo Scientific) as standard. The amount of BpMHAO HAO-like domain in both EcHAO and SoHAO was evaluated based on the western blot band intensity (see below).

The copper content was determined using a colorimetric method with a similar approach as Nóbrega *et al* [76]. In brief, protein samples are incubated with 100 µM of hydrochloric acid and 9 % of trichloroacetic acid at room temperature and, after the precipitate is formed, follows a centrifugation at 11,000 g for 5 minutes. The supernatant, containing the free copper, was then incubated with 20 mM ascorbic acid for 10 min. A solution of 2,2-biquinoline, 0.5 mg/mL, was added and the absorbance was measured at 546 nm (ε_546nm_ = 6.3 mM ^-1^cm ^-1^). The copper content was also assessed by inductively coupled plasma atomic emission spectroscopy, ICP-AES, at Laboratório de Análises at FCT-NOVA.

### Western-Blot

Western blotting analysis was carried out following the specific protocol from ProteinTech for Anti-His-Tag Monoclonal Antibody. An SDS-PAGE was performed, where the samples were pre-heated at 95 °C for 10 minutes and then loaded into the gel. The gel ran for 15 minutes at 80 V and then for 1h 15 min at 180 V. At this point, the gel bands were transferred to a 0.45 µM nitrocellulose blotting membrane (Cytiva) in a transfer apparatus for 1 h at 100 V in a Tris/Glycine buffer. The membrane blotting was performed for 1h at room temperature with PBS-Tween and 5 % milk. Then, the membrane was washed with PBS-Tween and left at room temperature for 1h with an Anti-His-Tag Monoclonal Antibody (Proteintech) (1:1000 dilution). After, the membrane was washed three times with PBS-Tween for 10 minutes each and the results were visualized using iBright Imaging Systems (ThermoFisher) after incubation with ECL solution (Cytiva).

The quantification of the HAO-like domain in the cell extracts for both organisms followed the same protocol. In brief, different amounts of the BpMHAO MCo-like domain (0.01-0.05 µg) were used to build a calibration curve establishing the relation between amount of protein and western blot band intensity. This was then used to calculate the amount of HAO-like domain correspondent to the signal observed. At least three different western blot gels were performed for each cell extract (three different amounts applied in each gel) [77].

### UV-Visible and Electron Paramagnetic Resonance spectroscopies

The UV-visible spectra under aerobic conditions were obtained using a Shimadzu UV-1700 spectrophotometer, whereas in anaerobiosis were obtained using a Shimadzu UV-1800 spectrophotometer inside an anaerobic chamber (Coy Lab Products).

Electron Paramagnetic Resonance (EPR) spectroscopy was performed using a Bruker EMX spectrometer equipped with an Oxford Instruments ESR-900 continuous flow helium cryostat. MCo-like domain samples (100 µM) were prepared in 50 mM MOPS buffer at pH 7.5. Spectral simulations and image preparation were performed using SpinCount [78].

### Determination of the reduction potentials

The MCo-like domain reduction potentials were determined by UV-visible and EPR monitored spectroscopies. Both titrations were performed inside the anaerobic chamber. Protein sample, at 100 µM, was prepared in 50 mM Tris–HCl, pH 7.5 and anaerobically titrated by successive additions of a buffered sodium dithionite solution at pH 9 (50 mM Tris-HCl pH 9), in the presence of an O_2_ scavenging system (1 M of glucose, 4.6 mM of glucose oxidase and 92 nM of catalase) and a mixture of redox mediators (1 µM of each in the assay): Potassium ferricyanide (E’_0_ = +430 mV), N,N dimethyl-p-phenylenediamine (E’_0_ = +430mV), 1,2- naphtoquinone-4-sulphonic acid (E’_0_ = +215 mV), 1,2 naphotoquinone (E’_0_ = +180 mV), trimethylhydroquinone (E’_0_ = +115 mV), phenazine methosulfate (E’_0_ = +80 mV), phenazine ethosulfate (E’_0_ = +55 mV), 5-hydroxy-1.4-naphthoquinone (E’_0_ = +30 mV), duroquinone (E’_0_ = +5 mV), menadione (E’_0_ −80 mV). A combined platinum electrode, with Ag/AgCl in 3.5 M KCl as a reference electrode, was used and calibrated at 20 °C against a saturated quinhydrone solution at pH 7. All reduction potentials were calculated in relation to the standard hydrogen electrode. The experimental data was manually adjusted with a Nernst equation for two consecutive monoelectronic transitions (Cu^2+^Cu^2+^ → Cu^2+^Cu^1+^ → Cu^1+^Cu^1+^).

A redox titration was also monitored by EPR spectroscopy. The MCo-like domain sample was prepared at 100 µM, under anaerobic conditions in 50 mM MOPS buffer at pH 7.5 and in the presence of the same O_2_ scavenging system. This sample were titrated by consecutive additions of a buffered sodium dithionite solution at pH 9 (50 mM Tris-HCl pH 9) in the presence of a redox mediators mixture containing the same mediators (0.7 mM each) used for the UV-visible titration plus: Plumbagin (E’0 = −40mV), Indigo trisulphonate (E’0 = −70 mV), Indigo disulphonate (E’0 = −110 mV), Phenazine (E’0 = −125 mV). The electrode and the respective calibration were performed as for the UV-visible monitored titration.

### Enzymatic activity assays

#### Laccase activity

The laccase activity was measured aerobically using ABTS (2,2’-azino-bis (3-ethylbenzothiazoline-6-sulfonic acid) (Alpha Aesar) or catechol (Aldrich) as substrates and measuring the absorbance at 420 nm (ε_420nm_ = 36 mM ^-1^.cm ^-1^) or 395 nm (ε_395nm_ = 1.4 mM^-1^.cm ^-1^) over time, following the oxidation of ABTS or catechol, respectively [79,80].The reaction was performed in different buffers: 100 mM glycine buffer at pH 3 and a 50 mM Tris-HCl pH 7.5 at 25 °C. The reactions were initiated by the addition of 2 µM of BpMHAO MCo-like domain and the rates were measured after the addition of the enzyme to the reaction mixture.

#### Nitrite reductase activity

The nitrite reductase activity was measured anaerobically, inside the anaerobic chamber, following the oxidation of methyl viologen (Sigma-Aldrich) at 730 nm (ε_730nm_ = 2.13 mM^-1^.cm^-1^) with a protocol described in [81]. Methyl viologen was prepared in 50 mM Tris-HCl pH 7.5 buffer and reduced overnight using zinc granules (Sigma-Aldrich). The reaction mixture was prepared by the addition of enzyme, i.e. 2 µM of MCo-like domain or 0.5 mg/mL for the soluble extracts (EcHAO, SoHAO, EcControl and SoControl) in 50 mM Tris-HCl pH 7.5 buffer and 500 µM of reduced methyl viologen. The reaction was initiated by the addition of 1 mM of a nitrite solution. The reaction rates were measured after the addition of nitrite to the reaction mixture.

#### Ferroxidase activity

Ferroxidase activity was monitored at 315 nm (36), following the oxidation of Fe^+2^ to Fe^+3^ using FeCl_2_ as a substrate. The reaction mixture contained 5 µM of MCo-like domain in 100 mM MOPS buffer at pH 7.5 with 150 mM NaCl, and the reaction started by adding 200 µM of FeCl_2_.

For the iron titration assays, the initial solution contained 5 µM MCo-like domain in 100 mM MOPS buffer at pH 7.5 with 150 mM NaCl and successive additions of FeCl_2_ were performed up to 60 equivalents. After each iron addition, the reaction was left to incubate for 10 minutes in a stirred cuvette before the spectrum was recorded. Similar assays were performed in anaerobiosis as control.

The ferroxidase activity as also measured amperometrically with a Clark-type electrode selective for oxygen (Oxygraph-2K, Oroboros Instrument, Innsbruck, Austria). The assays were performed in 100 mM MOPS pH 7.5 with 150 mM NaCl, at 25°C in an air-equilibrated buffer (≈ 250 µM of O_2_). The assay mixture contained 5 µM of the MCo-like domain and was initiated by the addition of 50 µM FeCl_2_ prepared in 100 mM MOPS buffer at pH 7.5 with 150 mM NaCl. Further additions were made after a stable oxygen measurement was achieved. The O_2_ consumed in each addition was calculated by the difference between the respective final and the initial O_2_ level and was used to calculate the Fe:O_2_ stoichiometry.

#### Hydroxylamine oxidase activity

The hydroxylamine oxidase activity was measured anaerobically following horse heart cytochrome *c* (Sigma) reduction at 550 nm (ε_550nm_ = 29.0 mM^-1^.cm ^-1^) (11) inside an anaerobic chamber. The reaction was initiated by the addition of 0.5 mg/mL of EcHAO or SoHAO to a reaction mixture containing 30 µM oxidized horse heart cytochrome *c* and 1 mM hydroxylamine in 50 mM Tris-HCl, pH 7.5. Control experiments were performed in a similar fashion but with EcControl and SoControl.

#### Hydroxylamine Reductase Activity

The hydroxylamine reductase activity was measured anaerobically by monitoring the oxidation of methyl viologen (Sigma-Aldrich) at 730 nm (ε_730nm_ = 2.13 mM^-1^.cm^-1^) (59). A methyl viologen stock solution was prepared as described above. The reaction mixture contained 500 µM of methyl viologen in 50 mM Tris-HCl pH 7.5 buffer, to which different amounts of hydroxylamine were added. The reaction started by the addition of the cell extracts (EcHAO, SoHAO and respective controls). The reaction rates were measured after the addition of the extracts. The assays where the stoichiometry between methyl viologen and hydroxylamine was measured were performed in a similar fashion, but with the successive addition of 100 µM of hydroxylamine in the same assay.

