## SupplementalMaterial for "A new player in the biochemistry of Anammox bacteria: a multidomain HAO-like protein"

### Supplementary Information

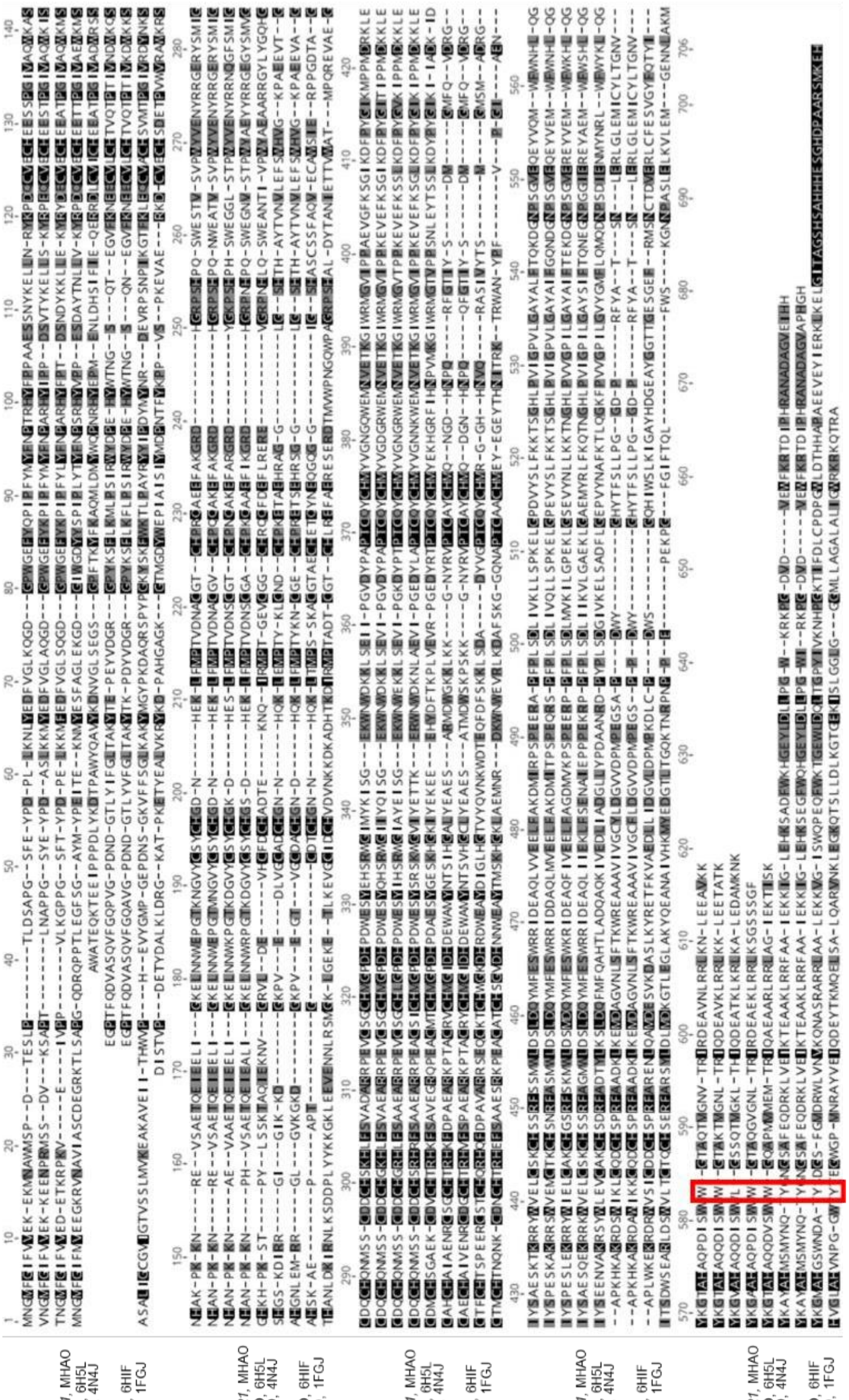

*Figure S1 – Amino acids sequence alignment of the HAO domain from the BpMHAO protein of Kuenenia stuttgartiensis, Brocadia pituitae and Jettenia caeni, and the HAO enzymes with known crystallographic structure, Kuenenia stuttgartiensis, Brocadia fulgida and Nitrosomonas europaea. The red box marks the tyrosine ligand of catalytic heme 4, which in BpMHAO is replaced by a tryptophan.*

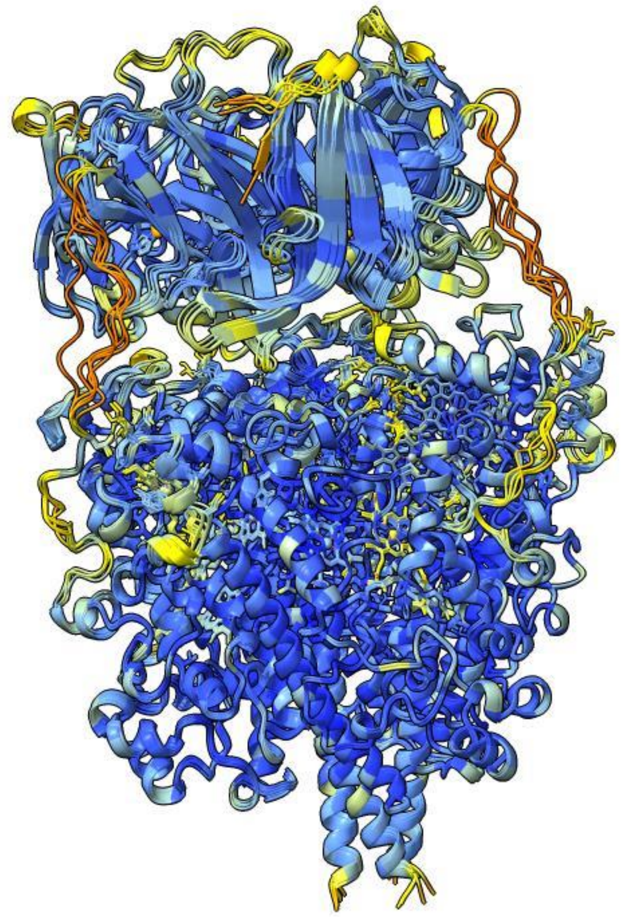

*Figure S2 - Superimposed AlphaFold3-predicted five models of the trimeric BpMHAO and coloured by per-residue pLDDT confidence scores. Blue indicates very high confidence (pLDDT >90), ligh blue indicates confident (90 > pLDDT > 70), yellow low confidence (70 > pLDDT >50), and orange indicates very low confidence (pLDDT <50).*

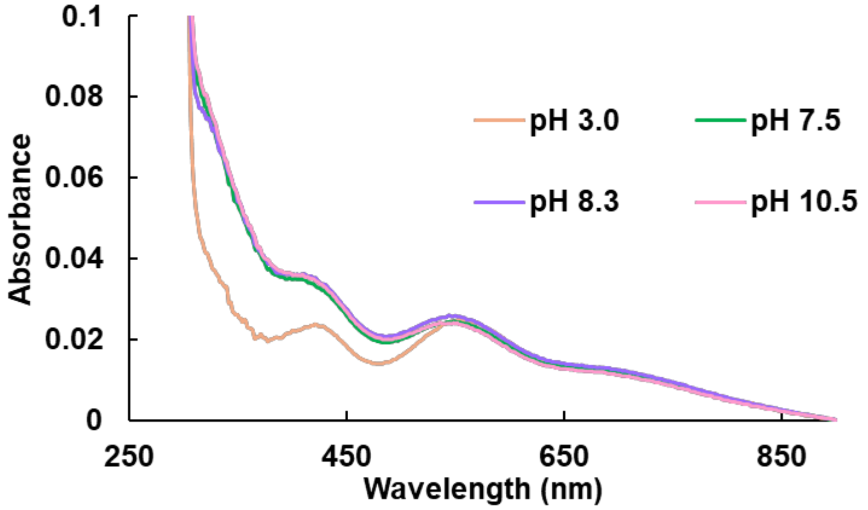


*Figure S3 - UV–visible spectra of 30 µM purified Mco-like domain recorded in buffers of varying pH. Spectrum at pH 3 were collected in 100 mM glycine (pH 3.0); spectrum at pH 7.5 in 50 mM Tris-HCl (pH 7.5); spectrum at pH 8.3 in 50 mM sodium phosphate (pH 8.3); and spectrum at pH 10.5 in 50 mM sodium phosphate (pH 10.5).*

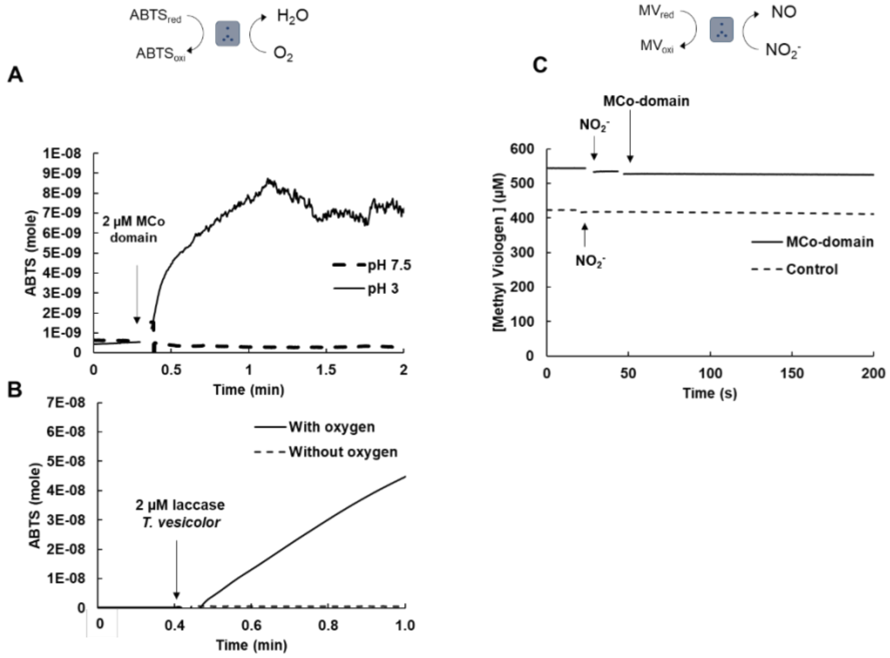


*Figure S4 - Laccase and nitrite reductase activities from the Mco-like domain. A) Laccase activity following the oxidation of the substrate ABTS 420 nm (ε = 36 mM-1 cm -1) aerobically with 2 μM Mco-domain in 50 mM Tris-HCl pH7.5 or 100 mM glycine pH 3. B) Laccase activity of the 2 μM small laccase Trametes versicolor activity following the oxidation of the substrate ABTS 420 nm (ε = 36 mM-1 cm -1) aerobically (black line) and anaerobically (dashed black line), in 50 mM sodium phosphate pH 6.0. C) Nitrite reductase activity of 2 μM of MCo domain, in 50 mM Tris-HCl pH 7.5, following the oxidation of methyl viologen at 730 nm (ε = 2.13 mM-1 cm-1) inside an anaerobic chamber. The assays from A), B) and C) are representative of three replicates.*

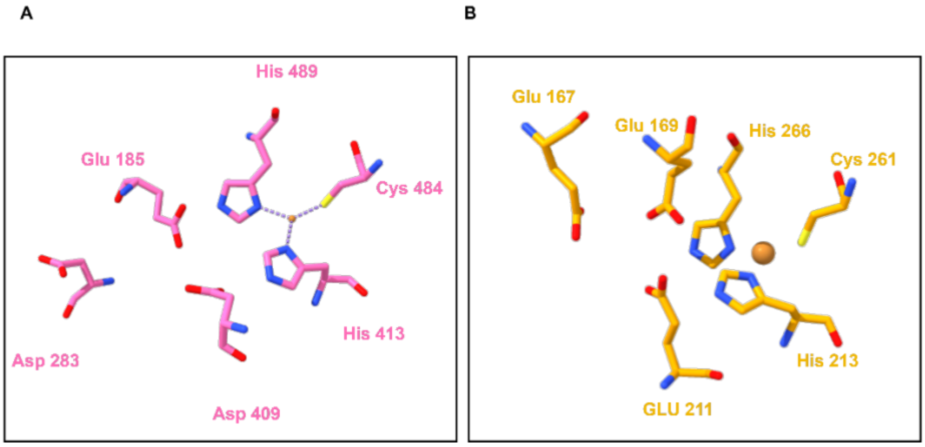


*Figure S5 - Close-up of the binding pockets of the T1 copper center of Fet3p ferroxidase from S. cerevisiae (PDB: 1ZPU) in pink and the structural model from the Mco domain from the BpMHAO in orange. A) The copper ligands and the amino acids responsible for ferroxidase activity in Fet3p (E185, D283, D409). B) Putative amino acids possibly responsible for the ferroxidase activity of the MCo domain from the BpMHAO protein.*

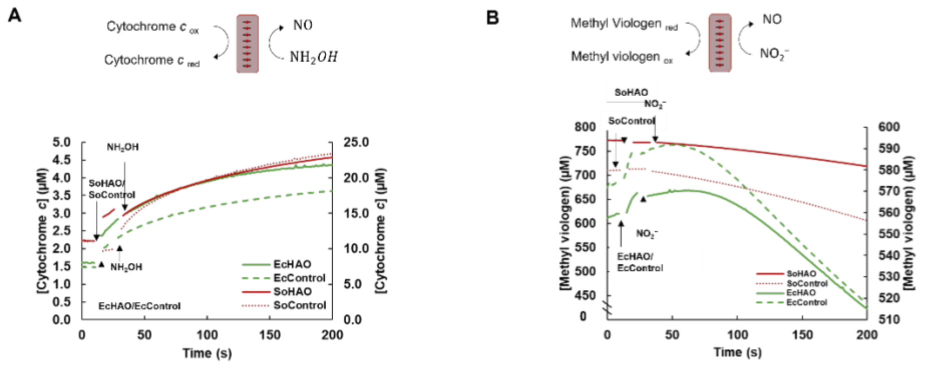


*Figure S6 - Hydroxylamine oxidase (A) and nitrite reductase (B) activities from EcHAO, EcControl, SoHAO and SoControl. A) Hydroxylamine oxidase activity following the reduction of cytochrome c at 550 nm (ε = 29 mM -1 cm -1), under anaerobic conditions for 0.5 mg.mL-1 of EcHAO (green line), EcControl (green dashed), SoHAO (red line), and SoControl (red dotted) of 1mM NH2OH in 50 mM Tris-HCl pH 7.5. Nitrite reductase activity of 0.5 mg. mL-1 of the same cell extracts with 1mM nitrite in 50 mM Tris-HCl pH 7.5, following the oxidation of methyl viologen at 730 nm (ε = 2.13 mM-1 cm-1) inside an anaerobic chamber. The assays from A) and B) are representative of three replicates.*
